# A Novel Role for Nucleolin in Splice Site Selection

**DOI:** 10.1101/2020.12.02.402354

**Authors:** Kinneret Shefer, Ayub Boulos, Valer Gotea, Yair Ben Chaim, Joseph Sperling, Laura Elnitski, Ruth Sperling

## Abstract

Latent 5’ splice sites are highly abundant in human introns, yet, are apparently not normally used. Splicing at most of these sites would incorporate in-frame stop codons generating nonsense mRNAs. Importantly, under stress and in cancer, splicing at latent sites is activated generating nonsense mRNAs from thousands of genes. Previous studies point to an unresolved RNA quality control mechanism that suppresses latent splicing independently of NMD. They further demonstrated a pivotal role for initiator-tRNA in this mechanism, through its interaction with the AUG codon, independent of its role in protein translation. To further elucidate this mechanism, here we searched for nuclear proteins directly bound to initiator-tRNA in the nucleus. We identified nucleolin (NCL), a multifunctional, abundant, and conserved protein, as a novel regulator of splice site selection. Starting with UV crosslinking, we show that NCL is directly and specifically interacting with initiator-tRNA in the nucleus, but not in the cytoplasm. In support of NCL involvement in this mechanism, we show activation of latent splicing in hundreds of transcripts upon NCL knockdown, disrupting gene transcripts involved in several important cellular pathways and cell metabolism functions (e.g. transcription factors, oncogenes, kinases, splicing factors, translation factors, and genes affecting cell motility, proliferation, and cellular trafficking). We thus propose NCL, a component of the endogenous spliceosome, through its direct interaction with initiator-tRNA and its effect on latent splicing as the first documented protein of a nuclear quality control mechanism that regulates splice site selection to protect cells from latent splicing that would generate defective mRNAs.

## Introduction

All multi-exon genes undergo regulated splicing, and most are subject to alternative splicing (AS), which provides a major source of diversity in the human proteome and contributes largely to regulation of gene expression (reviewed in refs. 1–6). Importantly, misregulation of splicing, through mutations or level changes of either regulatory signals or spliceosome components and splicing regulatory proteins, contributes to several human diseases, including cancer (7–10).

A key step in constitutive and alternative pre-mRNA splicing is recognition and selection of a consensus sequence that defines the 5’ splice site (5’SS). The 5’SS consensus sequence, AG/GTRAGT (where R denotes purine and “/” denotes the splice junction), is well defined and remarkably similar among many organisms, from mammals to plants (11–13). Surprisingly, surveys of human gene annotations demonstrated that intronic sequences abound in 5’SS consensus sequences that are not involved in splicing under normal conditions (termed latent 5’SS), exceeding the number of authentic 5’SSs by a factor of 6 to 9-fold (14, 15). Notably, the intronic sequences upstream of almost all (>98%) of these sites harbor at least one in-frame stop codon, and therefore have the potential to introduce premature termination codons (PTCs) into the alternatively spliced isoforms (15). mRNAs that contain PTCs are nonsense mRNAs that can be harmful to cells as their truncated proteins are likely non-functional and can have potentially deleterious dominant effects on the cell’s metabolism (16). Although splicing at latent sites has not been observed under normal growth, latent 5’SSs have been shown to be legitimate and can be activated. Latent splicing has been previously elicited in three different ways: (*i*) by eliminating the stop codons (in several gene constructs) either by point mutations that converted the stop codons to sense codons or by indels that shift the reading frame upstream of the stop codons (17, 18); (*ii*) by disrupting the reading frame through mutating the start ATG codon (19, 20); or (*iii*) by subjecting cells to stress conditions, such as heat shock or in cancer (15, 18, 19, 21, 22).

Two scenarios can account for why splicing at latent sites have not been observed under normal growth: (*i*) splicing at latent 5’SSs does occur, but an RNA surveillance mechanism, such as NMD (23–25), rapidly and efficiently degrades the nonsense mRNAs to a level below detection; or (*ii*) there is an unknown suppression mechanism of splicing at latent 5’SSs that are preceded by at least one stop codon in-frame with the upstream exon. Experiments have ruled out the first scenario of NMD (17–19, 22), or degradation by a yet unknown RNA degradation mechanism (20), while fitting the second scenario of latent splicing suppression. Latent splicing was shown to be regulated through the maintenance of an open reading frame (17–20, 22) and can be up-regulated under stress conditions (15, 21, 22). These discoveries suggest the existence of an RNA quality control mechanism — termed suppression of splicing (SOS) — whose proposed function is to suppress the use of latent 5’SSs that would generate nonsense RNA transcripts (26, 27). Support for nuclear recognition of a PTC-harboring pre-mRNA and suppression of splicing to prevent such transcripts has been observed in a number of studies (28–30), including a study that showed nuclear retention of unspliced PTC-harboring transcripts at their genomic loci (31). However, the mechanism of SOS quality control remains nebulous.

Previous experiments showed that SOS regulation requires an open reading frame (17, 18) and an initiation codon (19, 20), suggesting a role for the initiator-tRNA (ini-tRNA) in this regulation (20). Indeed, ini-tRNA, which was found associated with the endogenous spliceosome, was found to act as a pre-mRNA splicing regulator, independent of its role in translation, and was identified as a potential SOS factor (20). Mutations in the AUG translation initiation codon led to activation of latent splicing, but could be compensated for by expressing ini-tRNA constructs carrying complementary anticodon mutations, which suppressed latent splicing. This effect was specific to ini-tRNA, as elongator-tRNA, whether having complementary mutation to the mutated AUG or not, did not rescue SOS (20). Thus, ini-tRNA that recognizes the AUG sequence through base pairing is a pivotal element in the predicted SOS mechanism. Its interaction with the AUG sequence, probably in a complex with auxiliary proteins, would establish a register for the recognition of the reading frame required for SOS (20, 27, 32), which otherwise would not be discernible in the nucleus. Ini-tRNA presumably functions at the initial steps of SOS, conveying SOS components to the pre-mRNA.

Building on the previous discovery that ini-tRNA represents a key element in SOS, here we search for cellular components that directly interact with ini-tRNA in the nucleus to help decipher the SOS mechanism. We identify nucleolin (NCL), an abundant, highly conserved and multifunctional protein, which we previously reported as being associated with the endogenous spliceosome (33), as a new SOS factor. We show that NCL is directly and specifically interacting with ini-tRNA in the nucleus, but not in the cytoplasm. Furthermore, when we knocked down NCL we observed activation of latent splicing at hundreds of latent 5’SSs that introduce in-frame STOP codons, disrupting gene transcripts involved in several important cellular pathways and cell metabolism functions. These results suggest a novel role for NCL in splice site selection as a component of the SOS quality control mechanism proposed to protect cells from the use of latent 5’SSs that would generate mRNA molecules with PTCs.

## Results

### Search for Proteins Directly Associated with ini-tRNA in the Nucleus

To further explore the mechanism of SOS, we searched for factors directly associated with ini-tRNA only in the nucleus but not in the cytoplasm. In our first round of experiments, we injected ^32^P-labeled ini-tRNA into either the nucleus or cytoplasm of *Xenopus laevis* (*Xenopus*) oocytes, followed by UV crosslinking and RNase digestion (**Figure 1A**; see Materials and Methods for details). We found ini-tRNA directly associated with specific proteins in the nuclei as represented by two radioactive bands of apparent MW of ~60 and ~120 kDa (**Figure 1B**; lanes 5-8). Samples not treated by RNase revealed two major bands of apparent MW of 120 kDA and 170 kDA (see **Figure 1D**). The 60 and 120 kDa bands were not found in the cytoplasmic fraction samples (which only contained a ~26 kDa band) (**Figure 1B**, lane 1) or when the sample was not exposed to UV radiation (**Figure 1B**, lane 3).

**Figure 1.**
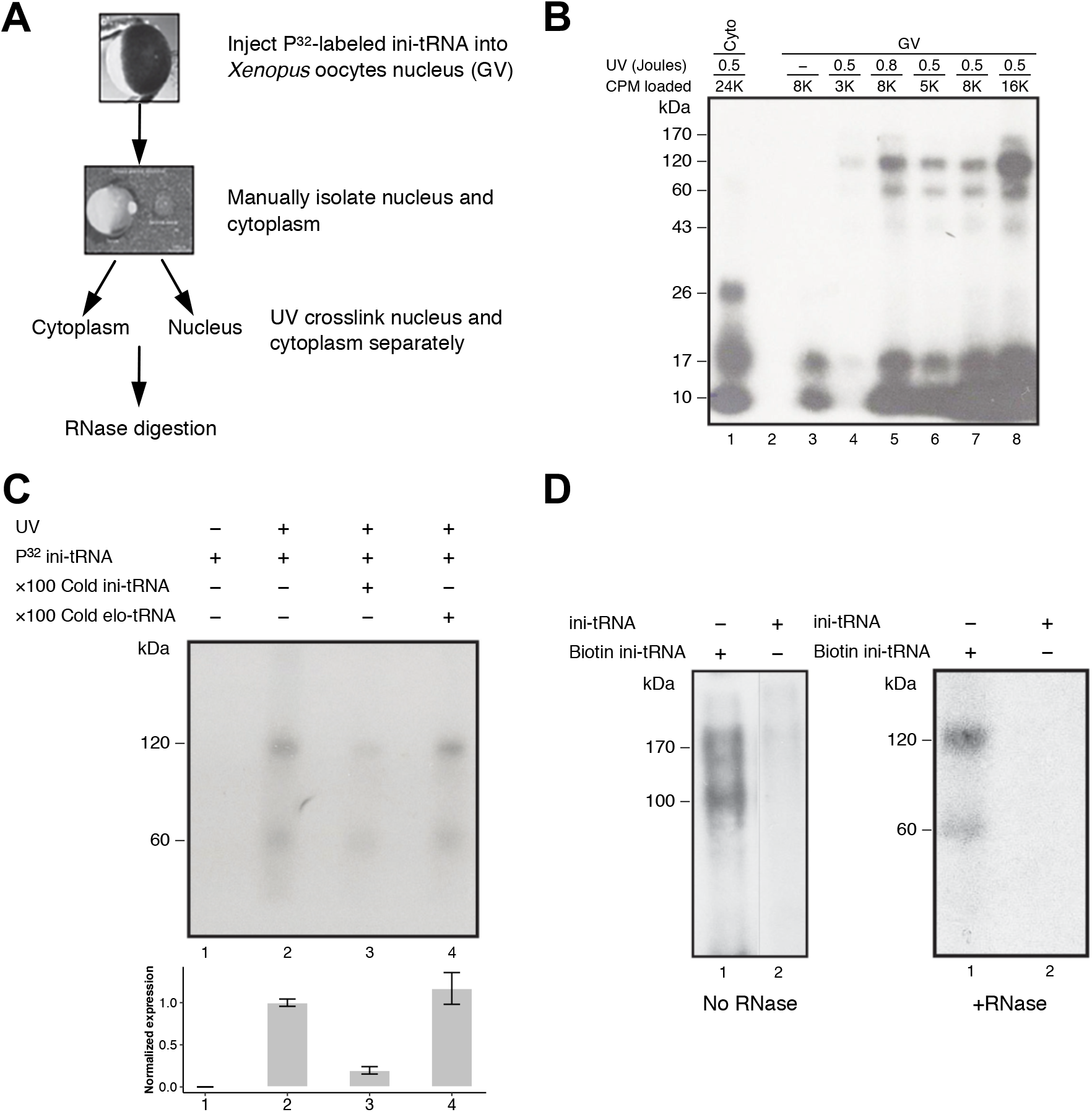
Specific crosslinking of proteins to ini-tRNA in the nucleus. (***A***) Scheme of the experiment. *In vitro* transcribed ^32^P-labeled ini-tRNA was injected into individual nuclei of *Xenopus* oocytes. Isolated nuclei and cytoplasm were crosslinked with UV light, and the crosslinked complexes were further digested with RNase. (***B***) SDS PAGE analysis of aliquots from experiments performed as described in *A*. Lane 1, UV crosslinked cytoplasm; lane 3, non-crosslinked nucleus; lanes 4-8, individual nuclei transfected with the indicated levels of ini-tRNA (cpm) and crosslinked by UV light at the indicated dose. (***C***) Specific association with ini-tRNA. The UV crosslinked bands are chased by 100X cold ini-tRNA, but not by 100X cold elongator-tRNA. Lower panel, quantification of the experiment (4 biological repeats for panels 1-3, and 3 biological repeats for panel 4) (***D***) Affinity purification of proteins bound to ini-tRNA. *In vitro* transcribed biotinylated ^32^P-labeled ini-tRNA was injected into individual nuclei of *Xenopus* oocytes [as described in (A)], UV crosslinked, and affinity purified on streptavidin magnetic beads and run on SDS PAGE. Control, non-biotinylated ^32^P-labeled ini-tRNA.

Importantly, this association of ini-tRNA with the factors represented by the 60 and 120 kDA bands is specific, because these bands, which only appeared after UV crosslinking and were not visible without it (**Figure 1C,** lanes 2 and 1, respectively), were chased by the addition of a hundred-fold cold ini-tRNA (**Figure 1C**, lane 3). Yet, they were hardly affected by a 100-fold excess cold elongator-tRNA (**Figure 1C**, lane 4). It should be pointed out that elongator-tRNA is a suitable negative control, because it is not associated with the endogenous spliceosome and it does not play a role in SOS (20). Specifically, activation of latent splicing induced by mutations in the translation initiation AUG codon, were suppressed by ini-tRNA constructs carrying anticodon mutations that compensate for the AUG mutations. In contrast, elongator-tRNA, either WT or mutated to complement the mutated AUG did not suppress activation of latent splicing, as demonstrated for three different transcripts (20). These observations indicate that factors with apparent molecular weight of 60 and 120 kDa were directly and specifically bound to ini-tRNA in the nucleus, but not in the cytoplasm.

To affinity purify the proteins associated with ini-tRNA in the nucleus, the experiment (see **Figure 1A**) was repeated, this time by injecting biotinylated ini-tRNA into *Xenopus* oocytes nuclei. Next, biotinylated ini-tRNAs with crosslinked proteins were affinity purified using streptavidin magnetic beads. As can be seen in **Figure 1D**, SDS PAGE of affinity-purified ini-tRNA with crosslinked components revealed two bands of associated components with apparent MW of 120 and 170 kDa, which yielded two respective bands of apparent MW of 60 and 120 kDa after RNase digestion (**Figure 1D**, lane 1, left and right, respectively). These two bands have the same apparent MW as the bands obtained previously, i.e., without affinity purification. The control (affinity-purified non-biotinylated ^32^P-labeled ini-tRNA) did not yield any product (**Figure 1D**, lane 2, left and right).

To increase the yield of affinity-purified peptides, we repeated the experiment, this time replacing injection by incubation of biotinylated ^32^P-labeled ini-tRNA with an extract from one isolated *Xenopus* nucleus or cytoplasm. Again, we obtained two bands of apparent MW of 170 and 120 kDa observed only in the nuclear extract and not in the cytoplasmic extract (**Figure 2A**, lanes 1 and 3, respectively) nor in the control (**Figure 2A**, lane 2). When we incubated our nuclear *Xenopus* extracts with increasing quantities (0.5–2 picomol) of biotinylated ^32^P-labeled ini-tRNA, the RNase treated products yielded 60 and 120 kDa bands again (**Figure 2B**, lanes 1-3). The intensity of the obtained bands increased concomitantly with increasing quantities of ini-tRNA. No bands were observed when the experiment was repeated with the cytoplasmic extract (**Figure 2B,** lanes 4-6), even with increased levels of ini-tRNA. Thus, the use of nuclear extract in our experiments enabled us to increase the amount of ini-tRNA from 10 fmol/nucleus to 2 picomol/nucleus in the procedure. These experiments show the direct and specific association between ini-tRNA and nuclear components having apparent MW of 60 kDa and 120 kDa.

**Figure 2.**
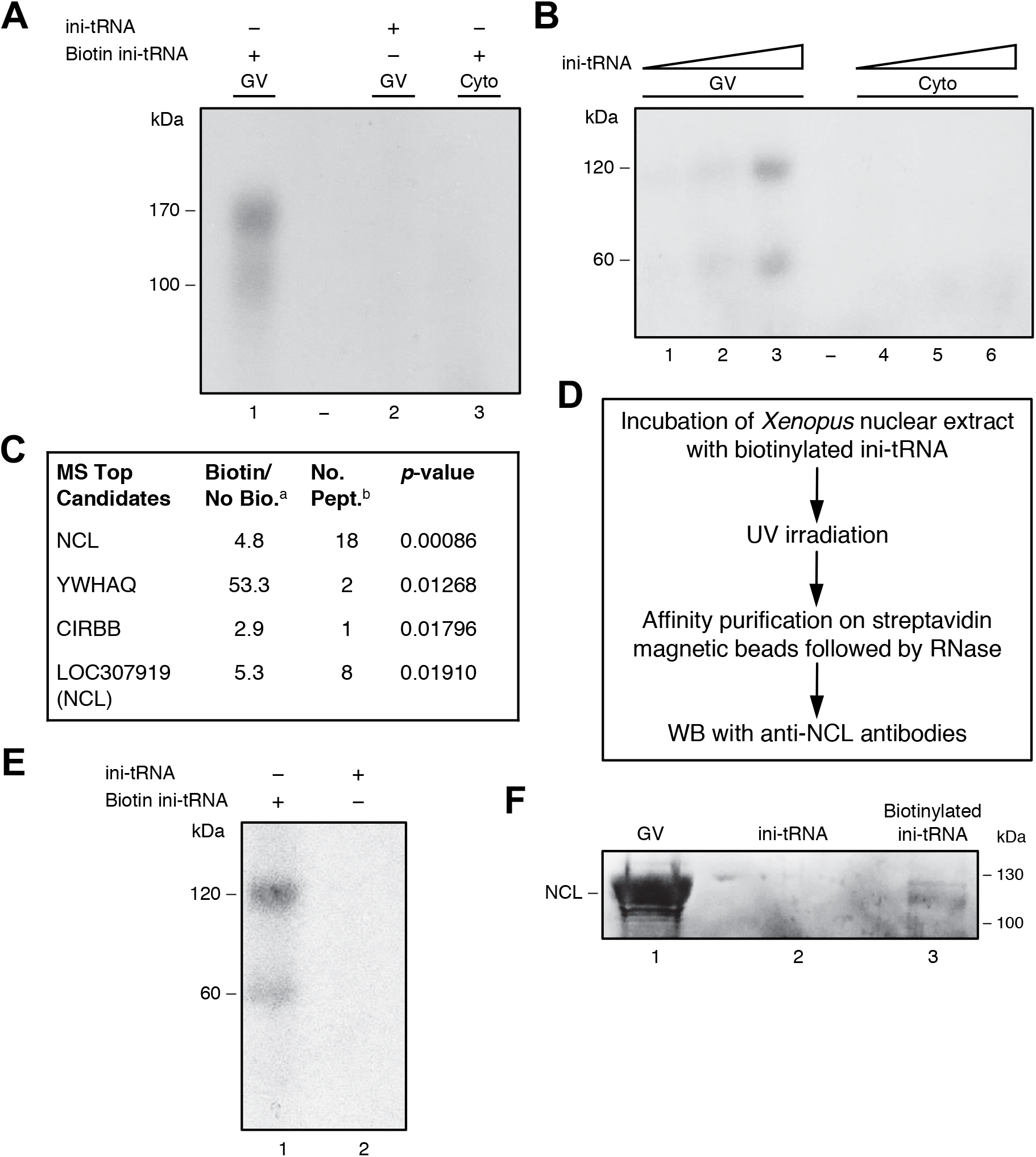
NCL is directly bound to ini-tRNA in the nucleus. (***A, B***) Proteins bound to ini-tRNA in nuclear extracts. Increasing quantities of *in vitro* transcribed biotinylated ^32^P-labeled ini-tRNA were incubated with nuclear (lanes 1-3) and cytoplasmic (lanes 4-6) extracts, UV crosslinked and affinity purified on streptavidin magnetic beads and run on SDS PAGE before (*A*) or after (*B*) RNase treatment. (***C***) Significant peptides associated with ini-tRNA in the nucleus based on mass spectrometry analysis; ^a^Bio/noBio indicates the fold change in biotin versus no biotin, average of three biological experiments; ^b^number of peptides (see also **Table S1**). (***D***) Scheme of the experiment described in *E*, *F* confirming NCL binding to ini-tRNA. (***E***) *In vitro* transcribed biotinylated ^32^P-labeled ini-tRNA was incubated with nuclear extract of *Xenopus* oocytes, UV crosslinked, and subjected to affinity purification on streptavidin magnetic beads, and run on SDS PAGE after RNase digestion (lane 1), and control non-biotinylated ^32^P-labeled ini-tRNA that went through the same procedure (lane 2). (***F***) Analogous experiment, followed by WB with anti-NCL antibodies. *Left*, GVs; *middle*, control non-biotinylated ini-tRNA; *right*, biotinylated ini-tRNA.

### Identification of Proteins Associated with ini-tRNA in the Nucleus Through Mass Spectrometry (MS) Analysis

Using the successful incubation and affinity purification approach described above, we performed a large-scale experiment, where we incubated nuclear extracts with non-radioactive biotinylated ini-tRNA (experimental samples) and non-biotinylated ini-tRNA (control). We also ran a parallel experiment under the same conditions but with biotinylated ^32^P-labeled ini-tRNA to help identify the relevant bands on the gel. We ran mass spectrometry (MS) analysis on the affinity-purified product (on bead digestion). **Figure 2C** summarizes the top results from three such biological repeats. To add stringency to our analysis, we considered genes as positive hits only when the ratio of biotin/no-biotin was larger than 3, with p-value <0.05, and when at least two peptides were identified (for the complete list see **Table S1**). According to the MS analysis, the top candidate protein was nucleolin (NCL). Our MS run identified eighteen of its peptides (*p*-value: 0.00086) and also peptides from the LOC397919 protein, another variant of NCL. Two additional MS experiments confirmed this result. In one of these two additional experiments, we extracted the proteins from the 60 kDa and 120 kDa bands and identified NCL in the 120 kDa band. It should be noted that NCL, with its acidic N-terminus, four RNA binding motifs in the middle, and a glycine/arginine-rich C-terminus, undergoes multiple post-translational modifications such as acetylation, ADP-ribosylation, glycosylation, methylation, and phosphorylation (34, 35). These modifications affect its migration on SDS gels, giving a band of apparent MW of 120 kDa (36). This altered migration is consistent with the migration of the upper band in our experiments (**Figures 1, 2**).

### Confirmation of NCL as a Candidate Protein Through Western Blot

To further validate the association of NCL of apparent MW of 120 kDA with ini-tRNA, we followed our UV crosslinking and affinity purification protocol (see **Figures 2D,E**) with western blot (WB) using anti-NCL antibodies (**Figure 2F**). We identified NCL in the 120 kDa band of untreated oocyte nuclei (germinal vesicle [GV] nuclei) (**Figure 2F**, lane 1), and in the 120 kDa band of affinity-purified proteins crosslinked to ini-tRNA (**Figure 2F**, lane 3), but not in the control (**Figure 2F**, lane 2). These results embolden the support for the identification of NCL as a protein that directly and specifically associates with ini-tRNA only in the nucleus of *Xenopus* oocytes.

### Annotating the Genomic Landscape of Latent 5’ Splice Sites

To assess whether NCL is affecting regulation of splicing at latent 5’ splice sites (here termed LSS), we first compiled a comprehensive annotation of these sites in the genome. We previously used a bioinformatics approach to identify potential LSSs in intronic sequences across the human genome, using data generated from expression exon microarrays (15). Using stringent criteria, we showed that LSSs are present in most human introns represented on the array (15). Here we used a more inclusive scoring threshold (MaxEntScan score (37) of 0 for intronic GT dinucleotides) to compile a comprehensive set of potential LSSs, filtered by known splice sites from multiple human gene annotation databases (see Materials and Methods for explanation, **Figures 3A,B**). This score threshold was chosen because it excluded the majority of intronic GT dinucleotides (less than 20% of intronic GTs score higher than 0) but was inclusive of potential new LSSs because more than 98.5% of annotated 5’SSs score higher than 0 (**Figure S1**). Limiting our search to LSSs located <1 kb downstream of annotated 5’SSs, we found a total of 1,381,214 LSSs (see Materials and Methods; **Table S2**). The majority of protein coding genes (nearly 89%) contained at least one LSS, with an average of nearly 80 LSSs per gene (median value: 55). The genes with the highest number of LSSs were *NBPF14* and *NOTCH2NLB*, which contained 1,190 LSSs each, whereas 72 other genes contained only one LSS. In general, we observed between 1 and 93 LSSs (median count: 8) within 1kb downstream of annotated 5’SSs, with multiple LSS occurrences being typical (in nearly 95% of cases).

**Figure 3.**
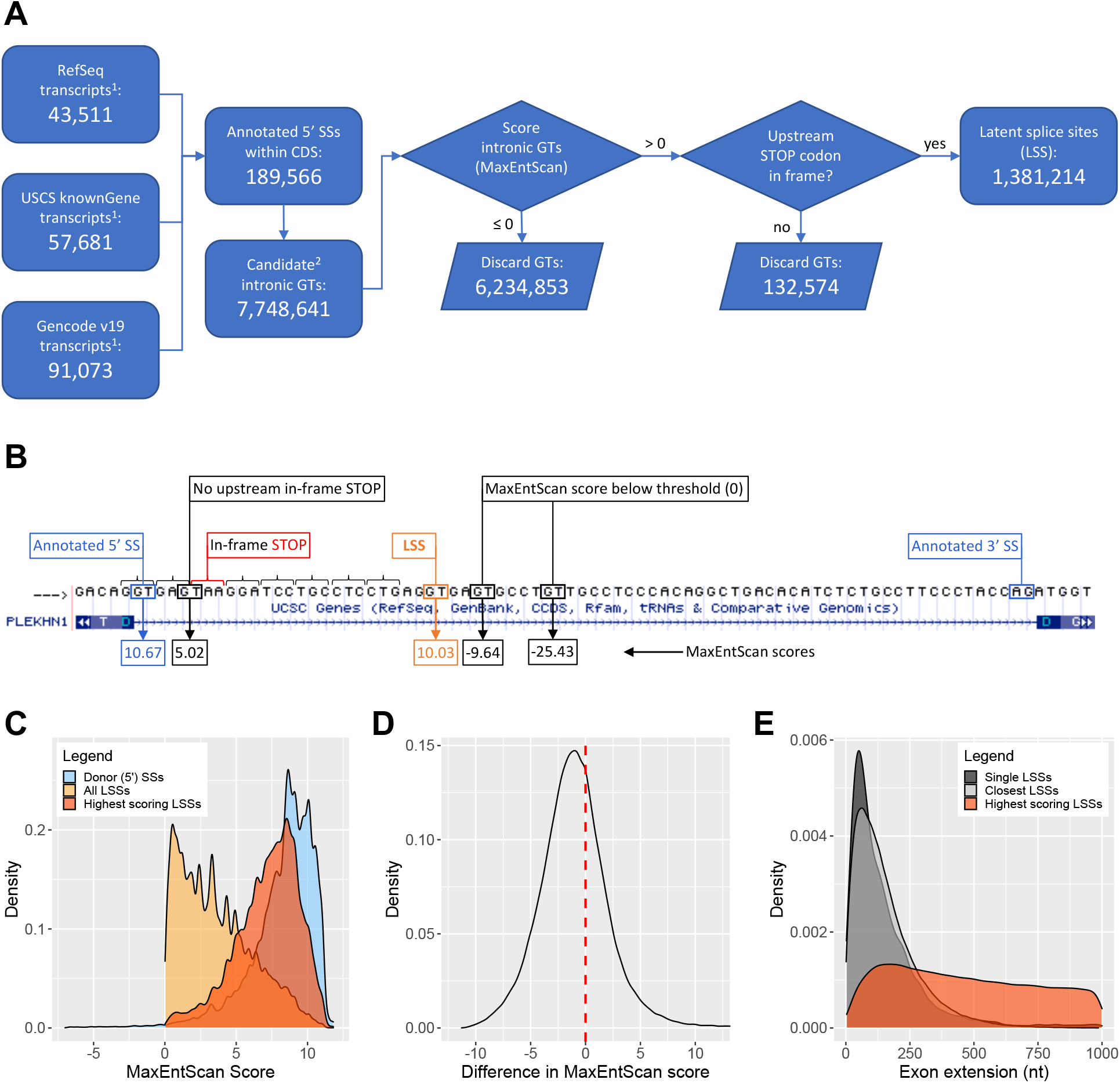
Latent splice sites (LSSs) in human genes. (***A***) Flowchart of computational steps performed to identify LSSs using the hg19 assembly of the human genome. ^1^Number of transcripts reflect only multiple-exon protein-coding transcripts annotated on chromsomes 1-22, X, and Y. ^2^Candidate GTs include only those GTs located in whitin-CDS intronic regions that would extend the upstream exon by at most 1000 nt while preserving a downstream intronic region of at least 20 nt (see also **Table S2**). (***B***) Examples of GTs evaluated for their potential LSS function in intron 15 of the *PLEKHN1* gene (RefSeq accession number NM_032129). (***C***) Distributions of MaxEntScan scores for LSSs and corresponding annotated donor 5’SSs. 0.9% of annotated 5’SSs have scores lower than −5 (not shown), and can be as low as −42.68. (***D***) Distribution of differences in MaxEntScan scores between the highest scoring LSSs located downstream of an annotated 5’SS and the score of that 5’SS. 34.2% of 5’SSs have at least one stronger LSS downstream of them. (***E***) Distributions of distances between LSSs and corresponding 5’SSs (i.e. exon extensions). “Single LSSs” denotes those cases where a single LSS can be found downstream of a specific 5’SS (8886 cases). For 5’SSs with more than one LSS downstream (159931 cases), distributions of exon extensions for both the closest and strongest LSSs are shown.

Most LSSs had lower MaxEntScan scores than the annotated 5’SSs upstream of them (median values of 3.19 and 8.68, respectively), and their score distribution was strongly skewed toward lower values (**Figure 3C**). However, if only the highest scoring LSSs downstream of each 5’SS was considered, the MaxEntScan score distribution was similar to the distribution of scores of annotated 5’SSs, albeit slightly shifted toward lower values (**Figure 3C**), highlighting the functional potential of intronic GT dinucleotides. For more than one third of established 5’SSs, there was at least one LSS downstream that scored higher (**Figure 3D**), highlighting the importance of the SOS mechanism for suppressing competition from these LSSs. The use of the highest scoring LSS instead of the annotated 5’SSs extended the upstream exon by an average of 469 bp (**Figure 3E**) and incorporated an in frame STOP codon. In all but 15.5% of cases with multiple LSS occurrences, the highest scoring LSS was preceded by LSSs with weaker MaxEntScan scores.

### NCL Knockdown by siRNA

To further explore the role of NCL in SOS, we knocked down NCL in HEK-293 cells using siRNA in two biological replicate samples (referred to as NCLsi samples), from which we isolated proteins and RNA (see Materials and Methods). Our controls were from two paired biological replicate samples treated with nontargeting siRNA (referred to as CONTsi), and untreated cells (referred to as CONT). Western blot analysis indicated that the NCL protein level was knocked down 25-fold (**Figure 4A**). HiSeq RNA sequencing (RNA-Seq) analyses yielded more than 230 million reads (126-bp, paired-end) for each sample, and about 84% of them were uniquely aligned to the hg19 assembly (**Table S3**). Standard differential expression analysis by DESeq2 revealed that NCL was the gene with the highest decrease in expression and the greatest fold change, as expected (**Table S4**). The normalized read counts for the NCL locus were reduced by more than 15-fold upon NCL knockdown (see **Figure 4B**; the DESeq2 model-based estimate of NCL downregulation is nearly 10-fold, FDR adjusted p = 3.5 × 10^−44^). Therefore, the WB and RNA-Seq analyses consistently indicated successful knockdown of NCL.

**Figure 4.**
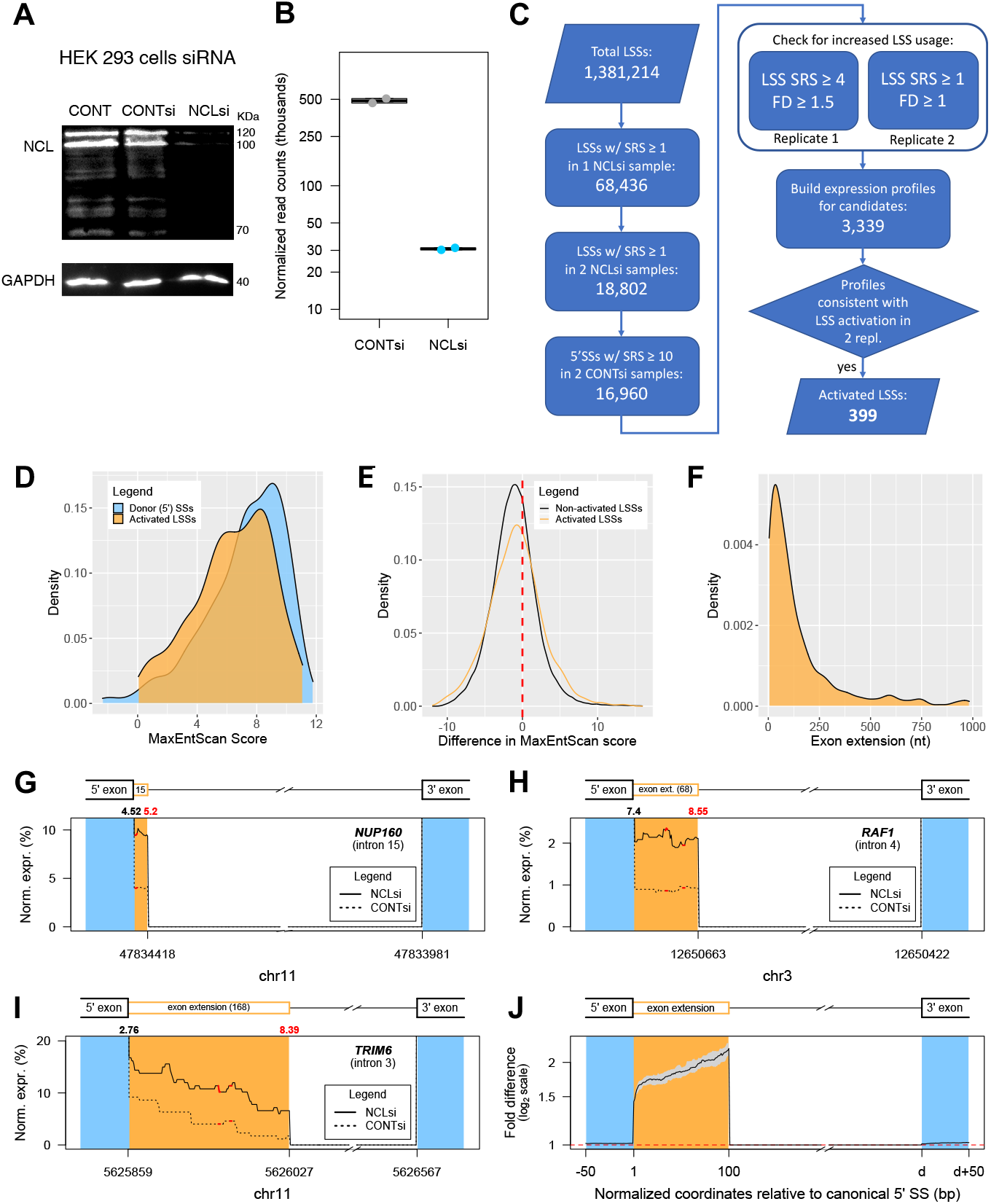
Identification of LSSs activated by knockdown of NCL. (***A*, *B***) HEK 293 cells were treated by siRNA against NCL (see Materials and Methods). As controls, we used non-targeting siRNA #1(Dharmacon), and non-treated cells. Proteins and RNA were extracted after 48 hr. (***A***) Proteins were analysed by SDS PAGE. (***B***) Analysis of NCL expression using RNA-Seq data (see Materials and Methods). Normalized reads of RNA from cells treated by siNCL (NCLsi) and siControl (CONTsi) (biological duplicates from each) are presented (see also **Table S4**). (***C***) The computational pipeline for identification of activated LSSs. Selection of LSS activated candidates was done based on the presence of split reads supporting the LSS junction. Replicate 1 and 2 can refer to either sample replicate pairs. Expression profiles for all candidates were analyzed individually to assure consistency between the two replicates. SRS – split read support (see also **Table S5**). (***D***) Distribution of MaxEntScan scores for the 399 activated LSS and the corresponding 385 annotated 5’SS upstream of them. (***E***) Distribution of differences in MaxEntScan scores between the activated LSSs and the corresponding upstream 5’SS (score_5’SS_ – score_LSS_). There are 401 unique LSS-5’SS pairs. Data for the set of non-activated LSSs was obtained with highest scoring LSSs downstream of 103,375 5’SSs with split read support of at least 10 reads for the canonical junction in both CONTsi samples. (***F***) Distribution of lengths for exon extensions caused by LSS activation. (***G-I***) Typical expression profiles of exon extension regions upstream of LSSs activated in NCLsi samples. Positions of in-frame STOPs are shown by red segments. Numbers on top of the graph indicate MxEntScan scores for the annotated 5’SS (black) and the LSS (red). The length of the exon extension is indicated in the corresponding segment of the gene model. (***J***) Composite profile showing the fold difference in expression level between NCLsi and CONTsi samples for all 399 cases of activated LSSs. The black line represents median fold difference values at nucleotide resolution, whereas the gray area represents 95% confidence interval for the median.

### NCL Knockdown Activates Latent Splicing

A detailed evaluation of the effect of NCL knockdown on splicing at LSSs was enabled by the depth of RNA sequencing. We used split reads (i.e., reads that align over two consecutive exons through an intron-induced gap) as the basic measure for LSS usage. These reads connect LSSs with the downstream acceptor SSs and represent direct evidence for the splicing of the isoform of interest. We found more than 35,000 LSSs in each sample being supported by at least one split read (1-NCLsi: 43679; 2-CONTsi: 35865; 7-NCLsi: 44124; 8-CONTsi: 41576). Overall, nearly 5% of all LSSs are supported by at least one split read in at least one of the NCLsi samples (**Figure 4C**). After adjusting for differences in the total number of mapped reads for each sample, we found in each replicate an excess of >4% (i.e. more than 1700) in the number of LSSs supported by split reads in NCLsi compared with CONTsi, supporting a role for NCL in regulating splicing at LSSs. To verify that NCL knockdown is linked to activation of splicing at many LSSs, we established a computational pipeline (**Figure 4C**) to separate cases of *bona fide* LSS activation from cases that are technical (e.g., spurious alignments) or biological (e.g., unannotated exons) artefacts. To define activated LSS, we required the NCLsi samples to exhibit increased usage of the LSS relative to the annotated upstream 5’SS, as well as higher levels of mapped reads throughout the exon extension region when compared to CONTsi samples (see Materials and Methods). We identified 399 LSSs in 362 genes that conform to all selection criteria (Table S5). The MaxEntScan score distribution for these 399 activated LSSs mimicked the score distribution for the 385 corresponding 5’SSs, with a slight shift toward smaller values (**Figure 4D**). Notably, 38.7% of activated LSSs showed higher or equal MaxEntScan scores to the corresponding upstream 5’SS (**Figures 4E**, **5B**). This finding, together with the similarity of the score distribution of activated LSSs to that of annotated 5’SSs, indicates that activated LSSs resemble genuine, annotated 5’SSs. Activated LSSs lead to exon extensions that range between 4 and 983 nt, with a median value of 81 nt (**Figure 4F**). Three cases of activated LSSs are illustrated in **Figures 4G-I**, and a median profile of the increased expression throughout the exon extension region for all 399 cases of activated LSSs is shown in **Figure 4J**.

**Figure 5.**
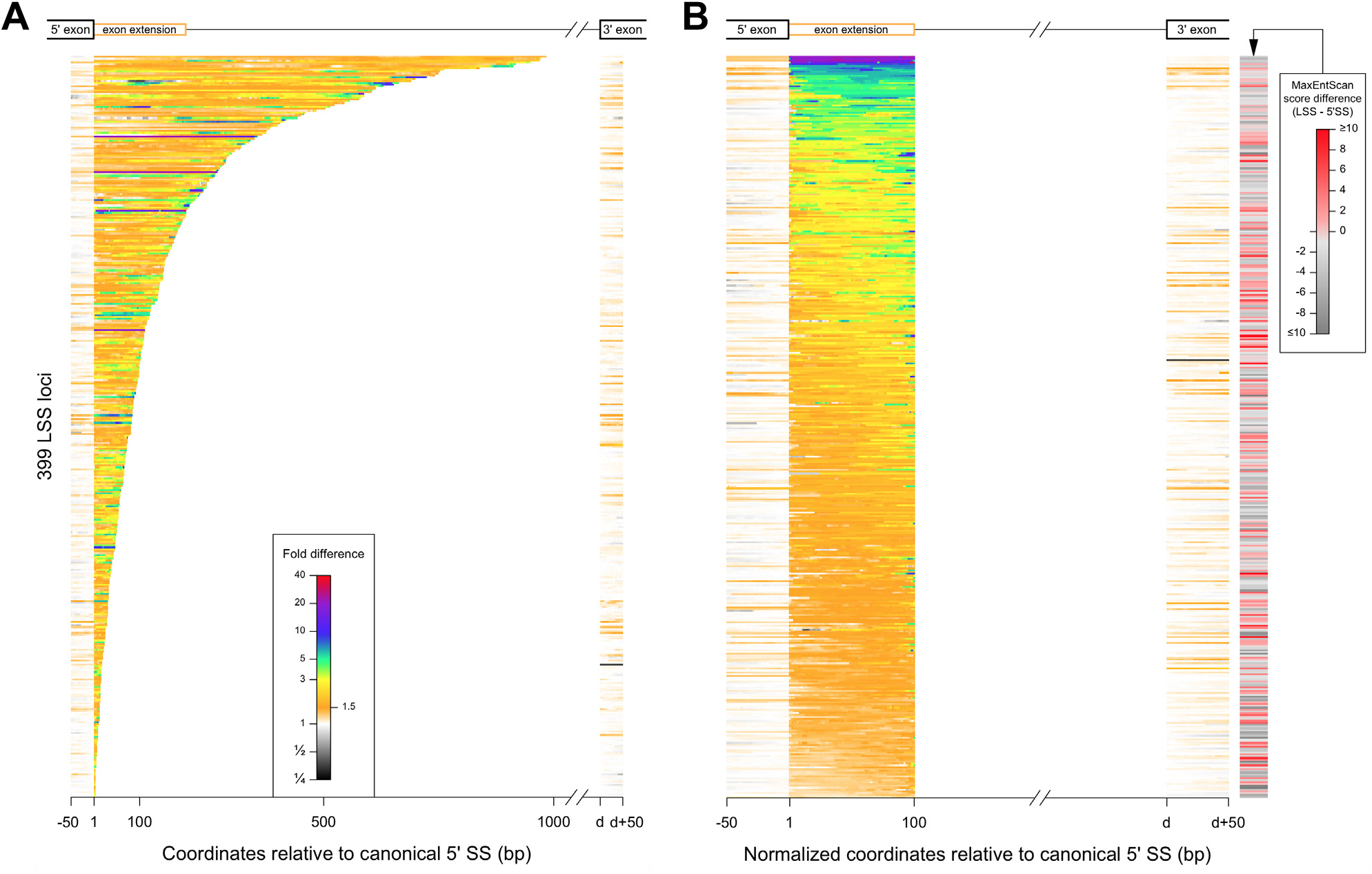
Overview of LSS activation upon NCL knockdown. Color-coded levels of fold-difference in expression levels is shown for all 399 activated LSSs. Fold difference was computed using normalized expression levels, where the normalization factor was the number of split reads supporting the corresponding annotated 5’SS (see also **Table S5**). (***A***) Data shown for actual length of exon extension regions, and LSSs are sorted based on the exon extension length. (***B***) LSSs are sorted based on the average fold difference across the exon extension region, which is shown as a standardized length of 100 bp. The difference in MaxEntScan score between the LSS and the corresponding 5’SS is shown on the right. Shades of red correspond to cases where the LSS has a score higher or close to the score of the corresponding 5’SS. The score is not significntly correlated with the increase in expression observed for the exon extension (Spearman’s *ρ* = −0.051, p = 0.3).

The heatmaps (**Figure 5**) provide a comprehensive view of the increase in expression associated with all activated LSSs upon NCL knock down, illustrating the wide range of length distribution of activated latent exons (**Figure 5A**). **Figure 5B** displays the average fold difference of activation of latent exons upon NCL knock down, each standardized to a length of 100 nt. It also portrays the MaxEntScan score difference between the activated LSS and the corresponding upstream 5’SS. Four activated LSSs are located in the *NCL* gene itself: three in intron 9 and one in intron 10. In **Figure 5B**, these four LSSs appear at the top of the graph, as they show the highest increase in expression levels of the exon extension regions.

We noted that LSS activation can be detected in isoforms with a wide range of expression levels. In a small number of cases, we observed increased usage (i.e., activation) of the same LSS within the context of different transcripts (i.e., the LSS is spliced with two different downstream 3’SSs, or with two different upstream 5’SSs). For example, an LSS located in intron 4 of *HMGN1* (chr21:40719840) appeared activated in two isoforms whose expression levels differ by almost 30-fold between the isoforms.

### Gene Transcripts Affected by NCL Knockdown

The cases with the strongest split read support for LSS activation after NCL knockdown are illustrated in **Figure 6A** (see also **Table S5**). This list includes proteins involved in several important cellular pathways and cell metabolism functions, such as transcription factors, oncogenes, kinases, splicing factors, translation factors, genes affecting cell motility, proliferation, and cellular trafficking, as well as NCL. These important functions can be found associated with many of the 362 genes affected by LSS activation, in agreement with the SOS being a general quality control mechanism.

**Figure 6.**
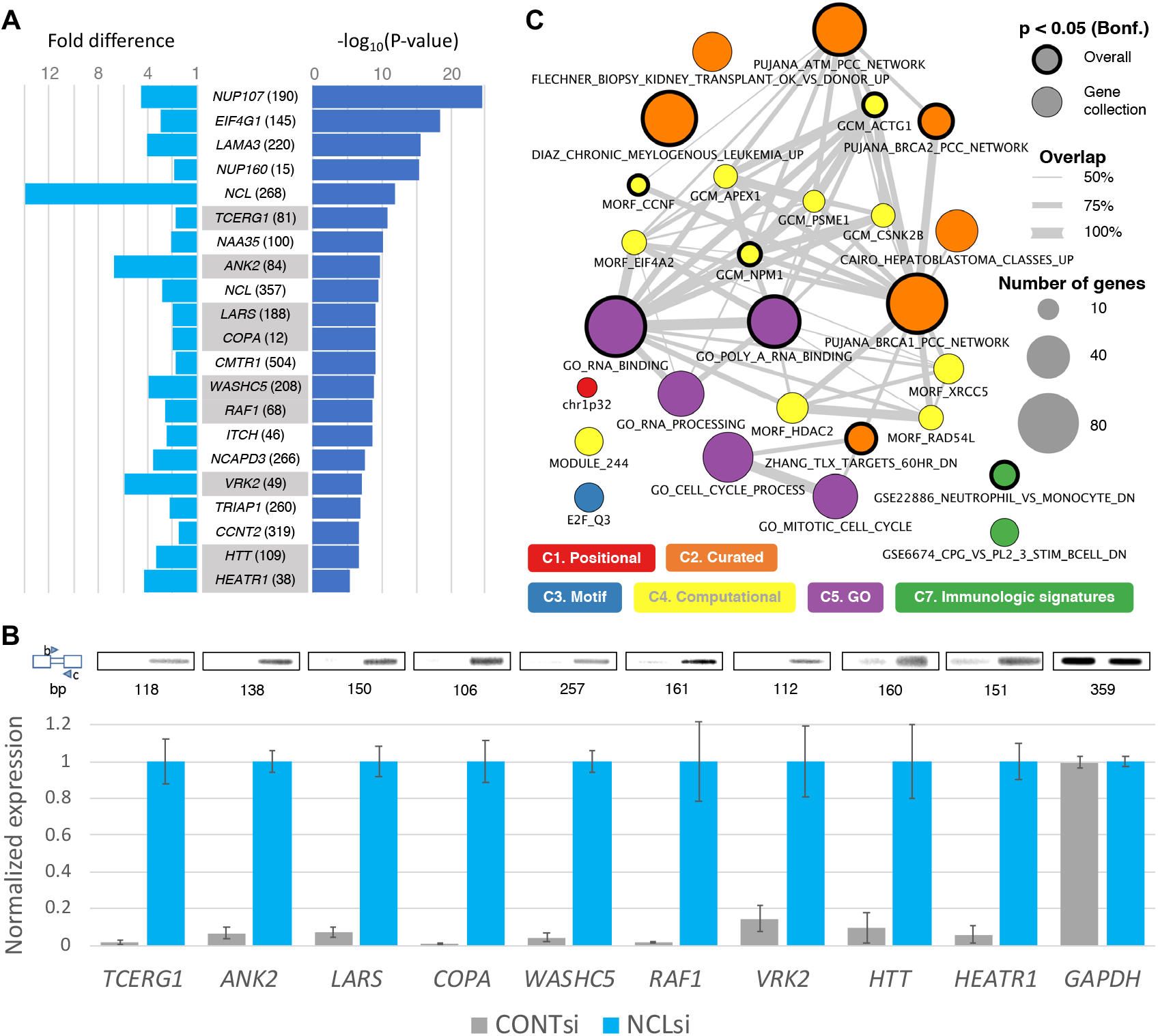
Genes with activated LSSs upon NCL knockdown. (***A***) List of 20 activated LSSs with strongest support based on split reads from the RNA-Seq experiments. Cases are ranked based on the p-value obtained using one-sided Fisher’s exact test with the number of reads supporting the LSS and the annotates 5’SS in the NCLsi and CTRLsi samples (values of the two replicates were combined using Fisher’s method). The fold difference in normalized LSS usage between NCLsi and CTRLsi samples is shown in light blue (the value shown corresponds to the geometric mean between the two replicates). Name of the gene and length of the exon extension are also provided (complete details for these cases can be found in **Table S5**). Cases experimentally validated through RT-PCR are highlighted. The case in the *HEATR1* gene was ranked 32nd, but is included here to show it was also experimatally validated. (***B***) RT-PCR validation of activation of latent splicing at LSSs in nine genes expressed in NCLsi treated HEK 293T cells. CONTsi treated cell were used as control. Numbers below PCR bands represent the sizes of the PCR products obtained with primers designed to match the schematic representation shown on the left (boxes represent exons, narrow box represents latent exon, i.e. exon extension, triangles represent primers). Bars represent averages and SEMs for three biological replicates. *TCERG1*, Transcription Elongation Regulator 1; *ANK2*, Ankyrin 2; *LARS*, Leucyl-tRNA Synthetase; *COPA*, Coatomer Protein Complex Subunit Alpha; *WASHC5*, WASH complex subunit 5; *RAF1*, Proto-Oncogene, Serine/Threonine Kinase; *VRK2*, vaccinia-related kinase (VRK) of serine/threonine kinase 2; *HTT*, Huntingtin; *HEATR1*, HEAT Repeat Containing 1. GAPDH, glyceraldehyde-3-phosphate dehydrogenase was used for normalization. All PCR products were verified by sequencing. (***C***) Gene sets from MSigDB over-represented among the 362 genes with activated LSSs. Circle areas correspond to the number of genes. Gray lines connect gene sets that share at least 50% of the genes in the smaller set. Thick circle borders correspond to gene sets that remain significant after stringent overall Bonferroni correction for multiple testing (17810 total tests) (see also **Table S6**).

For validation of the RNA-Seq results through an independent method, we selected nine gene transcripts (*ANK2, TCERG1, LARS, WASHC5, COPA, RAF1, VRK2, HEATR1*, and *HTT*) that are among the cases with the strongest RNA-Seq evidence for LSS activation (See **Figure 6A**). Using RT-PCR, we validated the RNA-Seq results showing that latent splicing was significantly elevated in response to knockdown of NCL as compared with the control cells (**Figure 6B**).

To learn whether LSS activation affects specific gene categories preferentially, we evaluated the over-representation of genes with activated LSSs among the gene sets included in all eight gene collections that make up the MSigDB database (i.e., numerous gene sets are within each collection). We found 27 gene sets with significant enrichment (after Bonferroni correction at gene collection level), 11 of which remained significant after stringent Bonferroni adjustment for the total number of 17,810 gene sets in MSigDB (**Figure 6C**, **Table S6**). The most significant (p = 5.9 × 10^−10^) and largest overlap (84 genes) observed was for genes that encode RNA binding proteins according to their gene ontology annotation. Subsets of these genes make up large fractions of many of the other 27 gene sets with significant enrichments (**Table S6**), which could explain the diversity of gene sets significantly affected by LSS activation. Moreover, given that many RNA binding proteins have essential cellular roles that include repair of DNA damage, DNA replication, chromatin regulation, transcription, splicing, translation, protein folding, and cell proliferation, these results suggest that the SOS mechanism plays a critical role in protecting the integrity of cellular functions.

### Further Evidence Supporting the Relevance of NCL in SOS

Our transcriptome survey through RNA-seq revealed increased latent splicing in hundreds of coding transcripts upon NCL knockdown. However, activation of latent splicing can be confounded and exacerbated by experimental stress (15, 26), such as the transfection procedure itself. To address this possibility, we switched sample labels between cases and controls and re-applied the computational pipeline to identify LSSs with increased usage. Under these conditions, we identified only 82 instances of LSSs activated in CONTsi compared to NCLsi samples (**Tables 1**, **S7**), nearly 5-fold fewer than identified with original sample labels. This finding indicates that NCL knockdown, rather than experimental stress, represents the main driver for the increase in latent splicing observed in NCLsi compared to control samples.

**Table 1.** NCL knockdown preferentially activates splicing at latent 5’SS that introduce PTCs.

| A5SS<sup>a</sup><br>types | Introduce<br>PTC | A5SS activated per sample type,<br>initial criteria<sup>b</sup> |  |  | A5SS activated per sample type, stringent<br>criteria<sup>c</sup> |  |  |
| --- | --- | --- | --- | --- | --- | --- | --- |
|  |  | NCLsi | CONTsi<sup>d</sup> | p-value<sup>e</sup> | NCLsi | CONTsi<sup>d</sup> | p-value<sup>e</sup> |
| LSS | Yes | 399 | 82 | $2.3 \times 10^{-11}$ | 187 | 23 | $4.2 \times 10^{-5}$ |
| adSS<sub>3n</sub> | No | 50 | 50 | N/A | 25 | 16 | N/A |
| adSS<sub>fs</sub> | Yes | 221 | 52 | $8.5 \times 10^{-9}$ | 112 | 20 | $1.6 \times 10^{-3}$ |

<sup>a</sup>Alternative 5' splice site. <sup>b</sup>Initial activation criteria require read support of at least 4 for the latent junction and increase in relative usage of at least 50% relative to the control sample in one replicate. For the second replicate, the latent junction is required to be supported by at least one read, while any increase in the relative usage is observed.
<sup>c</sup>Activation criteria (1.5-fold increase, read support of at least 4) are imposed to both replicates. <sup>d</sup>Activation of a 5SSs is determined by switching NCLsi and CONTsi sample labels and applying the pipeline described in Figure 4C.
<sup>e</sup>P-value was computed with one-sided Fisher's exact test against the adSS<sub>3n</sub> set.

To investigate whether the role of NCL is specifically linked to the presence of PTCs, we analyzed cases of latent splicing where no PTCs are incorporated. Specifically, these cases have no in-frame STOP codon in the exon extension region, and the length of the exon extension is a multiple a three, so as not to change the reading frame. We identified a total of 43,702 such potential alternative donor SSs, referred to as adSS_3n_ (**Table S8**). Using the methodology described in **Figure 4C** for LSSs, we identified 50 cases of activated adSS_3n_ in the NCLsi samples compared to controls (**Tables 1**, **S9**). Upon label switching, we also identified 50 instances (**Table S10**), suggesting that NCL knockdown has little impact on the choice of alternative donor SSs when no PTCs are introduced. Indeed, comparison of the LSS and adSS_3n_ activation reveals a significant bias toward LSS activation in NCLsi samples, i.e. upon NCL knockdown (**Table 1**). The same effect was observed even with more stringent criteria for activation of latent splicing (**Table 1**), supporting the hypothesis of NCL involvement in a mechanism that prevents generation of PTC-containing mRNAs.

To further investigate the relationship between NCL knockdown and the presence of PTCs, we also analyzed cases of latent splicing where PTCs are incorporated in downstream exons through a shift of the reading frame. For this purpose we searched for alternative donor SSs that do not introduce PTCs in the exon extension, but extend the exon by lengths that are not multiple of three, which would consequently cause a reading frame shift and insertion of a PTC downstream. We found a total of 98,349 such potential alternative donor SSs, referred to as adSSfs (**Table S11**). Analysis of RNA-seq data revealed 221 cases of activated adSSfs in NCLsi compared to control samples (**Tables 1, S12**), and 52 activated adSSfs sites were found upon label switching (**Tables 1**, **S13**). No significant difference can be observed between activation patterns of LSSs and adSSfs (p=0.28, initial criteria; p=0.17, stringent criteria, one-sided Fisher’s exact test), supporting a general role for NCL in suppressing PTC-inducing splicing events. In contrast, these data reveal a significant bias toward activation of adSS_fs_ compared to adSS_3n_ in NCLsi samples (i.e. upon NCL knockdown), similar to LSSs (**Table 1**). Therefore, our multiple complementary analyses support a general role of NCL in preventing PTC-harboring mRNAs.

## Discussion

### NCL is Directly Associated with ini-tRNA in the Nucleus

Previous studies showed that ini-tRNA, which is associated with the endogenous spliceosome, play a role in SOS (20). Here we identified NCL as directly and specifically associated with ini-tRNA in the nucleus, and not in the cytoplasm. Through our UV crosslinking experiments, we identified factors that interact directly and specifically with ini-tRNA in the nucleus. The specificity of interaction of ini-tRNA with the bound factors was demonstrated by chase experiments using a hundred-fold excess of cold ini-tRNA. However, elongator-tRNA, which does not play a role in SOS, and is not associated with the endogenous spliceosome (20), when used at the same concentration (100-fold excess of cold elongator-tRNA) did not affect the interaction of ini-tRNA with the bound factors (**Figure 1C**, lane 3 and 4, respectively). Affinity purification and mass spectrometry analyses of the crosslinked components revealed NCL as a protein directly and specifically associated with ini-tRNA in the nucleus but not in the cytoplasm. The association is specific as demonstrated by the competition experiments. Furthermore, it should pointed out that although NCL is an abundant protein, the level of NCL protein expression in *Xenopus* ooctyes stage IV-V, used here, is low (38, 39). The identification of NCL as bound to ini-tRNA is in agreement with earlier findings of the association of NCL with the endogenous spliceosome (33). These findings indicate a role for NCL in splicing regulation, which we extend to further define a role in the SOS mechanism.

### NCL – a Multifunction Protein with Novel Role in the Spliceosome

The results of our current study show that NCL has a previously undescribed role in splice site selection. NCL is known to perform multiple functions in numerous cell locations (nucleolus, cytoplasm, cell membrane and nucleoplasm) and to incur multiple post-translational modifications that affect its cellular location and function (34, 40). It is involved in the synthesis and maturation of ribosomes in the nucleolus, and while many of its functions in other cell components are described [e.g. a role in Pol II transcription, DNA repair, chromatin decondensation, and genome stability (34, 41, 42)], most of the molecular details of its assorted functions are not understood. Furthermore, NCL is known to play a role in cancer, where its overexpression affects cell survival, proliferation, and invasion (34, 42, 43). Our characterization of a previously unknown function for NCL is an important addition to the current body of knowledge of a protein crucial to cell function and involved in an important and widespread disease, cancer.

Recent studies have indicated a connection of NCL with splicing factors (44–47). We have previously found NCL as an integral component of the endogenous spliceosome (supraspliceosome), where it was identified associated with specific supraspliceosomes at all splicing stages (33). Furthermore, NCL was also identified to be associated with the general population of supraspliceosomes (48). While these published reports did not define a specific role for NCL in splicing regulation, as we do here, all of their findings support our discovery of NCL’s role in splicing.

### NCL Proposed as a Novel Regulator of Splice Site Selection to Protect Cells from Insertion of PTC

Our RNA-seq analysis revealed activation of 399 LSSs when NCL was downregulated. It should be pointed out that the number of activated LSSs uncovered here is likely a conservative estimate, because all the latent mRNAs are PTC-bearing and would likely become substrates for downregulation by NMD, as was previously demonstrated (15), limiting their detection by RNA-seq. Notably, we were able to further validate a subset of 9/9 of these cases by RT-PCR (*TCERG1*, Transcription Elongation Regulator 1; *ANK2*, Ankyrin 2; *LARS*, Leucyl-tRNA Synthetase; *COPA*, Coatomer Protein Complex Subunit Alpha; *WASHC5*, WASH complex subunit 5; *RAF1*, Proto-Oncogene, Serine/Threonine Kinase; *VRK2*, vaccinia-related kinase (VRK) of serine/threonine kinase 2; *HTT*, Huntingtin; *HEATR1*, HEAT Repeat Containing 1). This validation provides support for our bioinformatic approach. Importantly, the MaxEntScan score distribution of activated LSSs closely resembles the distribution of genuine, annotated 5’SSs (**Figure 4D**). Furthermore, 38.7% of activated LSSs show higher or equal MaxEntScan scores to the corresponding upstream 5’SS (**Figures 4E, 5B**). For annotating potential LSSs we have used a rather relaxed MaxEntScan that covers 98.5% of 5’SSs. Notably, even when more stringent MaxEntScan score thresholds are imposed for LSS selection a large fraction of activated LSSs can be detected. For example, a very stringent score threshold of 6.77, encompassing 80% of annotated 5’SSs, allows us to detect 191 activated LSSs (48% of the activated LSSs). Even with this stringent criterion, we detect highly significant bias toward LSS activation compared to adSS3n (p = 7.2 × 10^−7^, one-sided Fisher’s exact test) upon NCL knockdown. These observations strongly support the involvement of NCL in a mechanism that selects splice sites and thereby prevents insertion of PTCs into mature transcripts.

The gene transcripts (see **Figure 6A** and **Table S5**) that underwent activation of latent splicing when NCL was downregulated encode for proteins that are involved in several important cellular pathways and cell metabolism functions; they include transcription factors, oncogenes, kinases, splicing factors, translation factors, and genes affecting cell motility, proliferation, and cellular trafficking, highlighting the importance of this regulation. This finding is consistent with previous studies both in magnitude and in functional diversity of affected genes (15). Interestingly, our gene set enrichment analysis revealed an enrichment of RNA binding proteins in many of the significant gene sets (**Figure 6C, Table S6**). On the one hand, this is surprising, because we show that LSSs are present in most genes, and NCL was identified as directly binding to ini-tRNA that is proposed to be a general component of the SOS mechanism that conveys the SOS components to the supraspliceosome (20). Also, a previous analysis of the effect of heat shock on SOS revealed activation of latent splicing in hundreds of gene transcripts, reflecting a wide repertoire of functional groups (15). On the other hand, this over-representation could be a reflection of the breadth of essential cellular functions performed by RNA binding proteins, and therefore, it is functionally critical that such genes are spliced properly. This is in line with recent observations that SS selection is tightly regulated in germ cells (49). Additional explanations for the observed enrichment in RNA binding proteins could include interaction with tissue-specific splicing factors or recruitment of splicing factors having high affinity for specific motifs within the pre-mRNA of RNA binding proteins. Yet, other explanations cannot be excluded at this stage. Further experiments directed at deciphering the molecular interactions are required to resolve this issue.

### Proposed Update of the SOS Speculative Model

The proposed SOS working model (20) includes the role of ini-tRNA as an SOS factor. It invokes however a highly speculative sense triplet-recognition mechanism that can be interrupted by stop codon-binding proteins. In this study, we identified NCL as a potential novel SOS factor by demonstrating that NCL is directly associated with ini-tRNA in the nucleus and its downregulation activates latent splicing at hundreds of gene transcripts. Using these results, we have updated our working model to include SOS as a quality control mechanism within the supraspliceosome that acts before splicing (**Figure 7**). The supraspliceosome is composed of four native spliceosomes, which are connected by the pre-mRNA (27, 50, 51). According to this model, first, splice site combinations are selected through the combinatorial interplay of positive and negative regulatory signals present in the pre-mRNA, which are recognized by trans-acting factors, resulting in the assembly of a supraspliceosome on each pre-mRNA (**Figure 7A**). Second, SOS endorses the right combinations of splice junctions (**Figure 7B**). The details of NCL role in SOS require further study.

**Figure 7.**
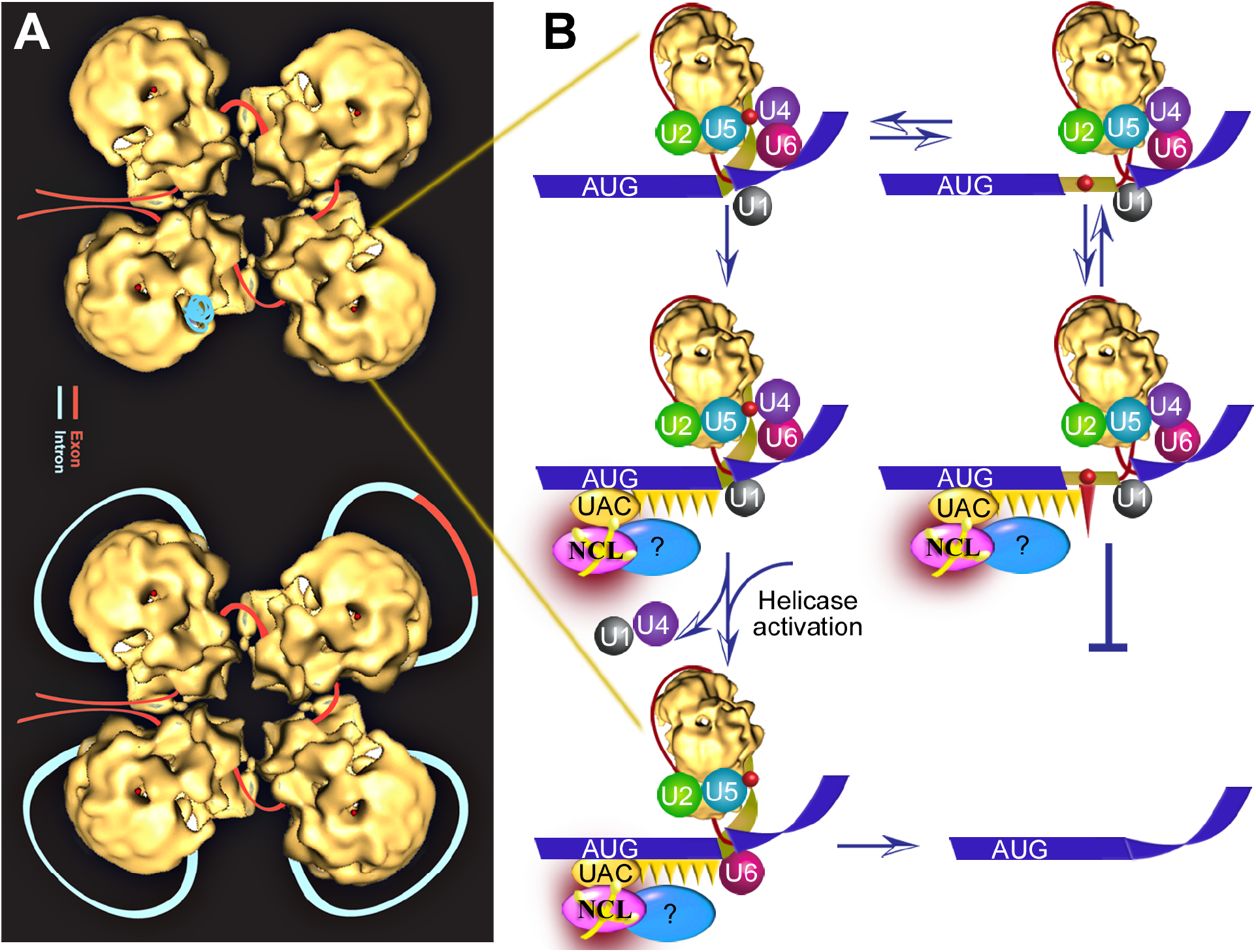
An updated speculative schematic model for the quality control function of SOS. (***A***) The supraspliceosome model (50, 51, 63–65). Exon, red; intron, light blue. (Top) The folded pre-mRNA that is not being processed is protected within the cavities of the native spliceosome. (Bottom) When a staining protocol that allows visualization of nucleic acids was used, RNA strands and loops were seen emanating from the supraspliceosomes (66). The RNA kept in the cavity likely unfolded and looped-out under these conditions. In the looped-out scheme an alternative exon is depicted in the upper right corner. **(*B***) Zoom into one spliceosome. Left scheme, splicing at the authentic 5**’**SS; right scheme, splicing at the latent 5**’**SS. Blue stripes, exons; red line, intron; yellow narrow stripe, latent exon; red circle, in-frame stop codon; circles, U snRNPs; orange ellipse (UAC), initiator-tRNA; purple ellipse, NCL directly bound with ini-tRNA; and blue ellipse, additional associated components; orange triangles, hypothesized triplet-binding proteins; red triangle, stop-codon-binding protein. Updated from ref. (20).

We proposed that the SOS mechanism is based on three elements. First, the AUG sequence is recognized by the complementary anticodon (UAC) of the ini-tRNA, which is in a complex with auxiliary proteins (20). NCL can now be added in the model as a protein directly binding ini-tRNA, likely along with additional proteins. This step helps to establish a register for the recognition of the reading frame. The second SOS element involves the cooperative polymerization of protein(s) that bind triplets of nucleotides and in the absence of a PTC it reaches the selected 5’SS; this step is the quality control “confirming” step of SS selection that triggers the remodeling of the spliceosome to its functional state (**Figure 7B**, left). The final SOS element is suppression of splicing in the presence of a PTC, perhaps through a competing interaction with a stop-codon-binding protein (e.g., a release factor-like protein). The unproductive complex may undergo a conformational change and revert to the productive splicing complex involving the authentic 5’SS, as indicated by the double arrows (**Figure 7B**, right). In cases where the quality control mechanism fails, for example in cases of stress, downstream mechanism/s (e.g., NMD in the cytoplasm) may engage to safeguard the robust control of the system.

## Conclusions

Our study identified NCL, a highly abundant and conserved protein, with multiple cellular functions involved in cancer, as a potential novel regulator of splice site selection and a component of the SOS mechanism that is proposed to protect cells from latent splicing that would generate transcripts with PTCs. We also identified novel splicing targets of NCL involved in multiple important cellular pathways and cell metabolism functions.

## Materials and Methods

### Model system

We chose the *Xenopus laevis* (*Xenopus*) oocyte system for our experiments because SOS appears to be evolutionarily conserved (22), and thus the high protein concentration in *Xenopus* oocyte nuclei could be used to identify the proteins interacting with ini-tRNA in the nucleus. Furthermore, this system has been successfully used as a model system to study numerous biological processes, including splicing (52, 53), and we previously used this system to show that injected ini-tRNA, which functions in SOS, is uncharged with an amino acid (20).

### *In vitro* Transcription of ini-tRNA

We generated a DNA template using pTRM, a plasmid carrying the WT ini-tRNA gene (54) and specific primers in a PCR reaction according to the manufacturer’s instructions, and as detailed in Supplementary Methods. The RNA was purified using the miRNeasy kit (217004 Qiagen), according to the manufacturer’s instructions, and analyzed on a denaturing gel.

### *In Vitro* Transcription of Elongator-tRNA

We annealed 1 mM of a primer having the elongator ini-tRNA antiparallel sequence to 1 mM T7 primer (90°C for 3 minutes), and cooled it to room temperature (RT; annealing buffer: 10 mM Tris-HCl pH 7.5, 50 mM NaCl). Then, we followed the transcription procedure explained above.

### Injection of ini-tRNA into the Nuclei of *Xenopus* Oocytes

We extracted *Xenopus* oocytes as previously described (55, 56), and as detailed in Supplementary Methods. A day after oocyte extraction, 10 fmol (20 nl) of ^32^P ini-tRNA was injected into the oocyte’s nucleus (Picospritzer, PLI-100; Medical Systems Corp.). After 30 minutes, nuclei were extracted manually, and nuclei were gently squeezed out into GV buffer (5 mM HEPES, 17 mM NaCl, 83 mM KCl). Each nucleus was gently washed and transferred to a new plate (57). Radioactivity of nuclei was measured (Perkin-Elmer Tri-Carbs 2900 TR Liquid Scintillation), and only nuclei having over half of the injected ini-tRNA were further assayed. Nuclei were irradiated at 4°C with an energy of 0.8 Joule (or 0.5 Joule as indicated) [Ultra-Lum UV Cross-Linker (UVC 515)]. Nuclei were digested by RNase A [1.75X10^−3^ RNase A units (Sigma R-6513) per 50 μl sample volume] and run on 8.7% SDS PAGE (1 nucleus/well).

### *In vitro* Binding of ini-tRNA to Nuclear Extract of *Xenopus* Oocytes

We incubated nuclear extract (one nucleus per reaction, as described above) with increasing quantities (0.5–2 picomol) of biotinylated ini-tRNA for 10 minutes and irradiated them with UV light (as described above).

### Affinity Purification of Biotinylated ini-tRNA and its Crosslinked Factors

We affinity purified ini-tRNA with crosslinked components using 5 μl (or else as indicated) C1 streptavidin magnetic beads (Invitrogen 65001) in B&W buffer according to the manufacturer’s protocol. The pellet was resuspended in SDS loading buffer, boiled at 90°C for 5 minutes, and run on an 8.7% SDS PAGE.

### RT-PCR Analysis

RT-PCR was performed on RNA extracted from the NCLsi and CONTsi treated cells as described (15), and detailed in Supplementary Methods, using the detailed sets of primers (see Supplementary Methods). The identity of all PCR products was confirmed by sequencing. Each experiment was repeated at least 3 times.

### Western blot (WB)

We ran our samples on 8.7% SDS PAGE and transferred them to a nylon membrane as described before (33). We performed WB analyses using anti-NCL antibodies (C23, cat # Sc-13057, Enco) and visualized our product with horseradish peroxidase conjugated to affinity-purified goat antirabbit IgG (1:5000 dilution; H + L; Jackson ImmunoResearch).

### Mass Spectrometry Analyses

We incubated 60 *Xenopus* oocyte nuclei with 120 picomol ini-tRNA (assay samples were prepared with biotinylated UTP and control samples were prepared with non-biotinylated UTP). All samples were affinity purified using 150 μl C1 streptavidin magnetic beads (as described above) and then resuspended in 100 μl 10 mM Tris buffer pH 7.5. We used in-solution, on-bead, tryptic digestion to prepare samples, as detailed in Supplementary Methods. Liquid Chromatography and Mass Spectrometry analyses are detailed in Supplementary Methods.

### Knockdown of NCL

We transfected siRNA targeted to NCL (L-003854-00-0005, ON-TARGETplus Human NCL (4691) siRNA-SMARTpool, Dharmacon), and non-targeting siRNA as control (D-001810-01-05 ON-TARGETplus Non-targeting siRNA #1, Dharmacon) into HEK 293 cells with TransIT-X2 (mc-MIR-6003, Mirus) according to the manufacturer’s instructions with some modifications. Cells grown in 6-well plates were transfected with 45 nM siRNA by using TransIT-X2 transfection reagent. After 24 h, medium was changed, and cells were transfected again, as before. After 48 h, total proteins and RNA were extracted (RNeasy Qiagen 74104) according to the manufacturer’s instructions and analyzed.

### RNA Library Preparation and Sequencing

Using the TruSeq® Stranded mRNA Sample Preparation kit (Illumina), we prepared a RNA library from four samples: (1)1-NCL-siRNA (1-NCLsi), (2)2-si-CONTROL (2-CONTsi), (3)7-NCL-siRNA (7-NCLsi), and (4)8-si-CONTROL (8-CONTsi). Briefly, polyA fraction (mRNA) was purified from 500 ng of total RNA for each sample, fragmented, and used to generate double stranded cDNA. We then performed end repair, A base addition, adapter ligation, and PCR amplification of the samples. We evaluated the libraries with Qubit and TapeStation. Sequencing libraries were constructed with barcodes to allow multiplexing of four samples on four sequencing lanes, which were then sequenced with the Illumina HiSeq 2500 V4 instrument. More than 234 million paired-end, 126-bp reads were generated in total for each sample (**Table S3**).

### Identification of Latent 5’SSs and Alternative Donor SSs

We compiled a comprehensive list of potential latent 5’ splice sites (LSSs) in the human genome by investigating all GT dinucleotides located in intronic regions of multi-exon coding transcripts. Upon passing a MaxEntScan score threshold of 0, the main criterion qualifying a GT dinucleotide as an LSS is the introduction of at least one in-frame STOP codon upstream (see Supplementary Methods for detailed information). Other alternative donor splice sites, such as adSS_3n_ (extend coding exons by lengths that are multiple of three) and adSSfs (extend coding exons by lengths that are not multiple of three), were selected only among GT dinucleotides that do not introduce upstream in-frame STOP codons.

### Analysis of RNA-Seq Data

We performed read alignment with the STAR (v2.3.1) aligner (58) against the hg19 (GRCh37) assembly of the human genome (for further details see Supplementary Methods). We used the QoRTs package (59) to quantify reads mapped to exon junctions and genes included in the Gencode v19 annotation track. Differential gene expression analysis was performed with the DESeq2 package (60) in R (v3.2.2) with two case samples (1-NCLsi and 7-NCLsi) and two controls (2-CONTsi, 8-CONTsi).

### Quantification of Increased LSS Usage

To compare LSS usage between samples we used the counts of reads mapped to the junction formed by the LSS and the downstream acceptor SS normalized by the counts of reads mapped to the corresponding canonical junction (i.e. # split-reads_LSS_ / # split-reads_5’SS_). As candidates for activation of latent splicing we considered those LSSs for which we observed at least one NCLsi replicate with at least 4 reads supporting the LSS and a fold increase ≥1.5 in normalized LSS usage in the NCLsi compared to the CONTsi sample. Additional conditions imposed to minimize the number of false positives are described in the Supplementary Methods.

### Analysis of Expression Profiles

To further verify that the observed increase in LSS usage corresponds to the extension of the upstream exon into the intronic region we built expression profiles by quantifying reads mapped throughout the exon extension region. We compared normalized NCLsi and CONTsi profiles and retained cases for which both replicates were consistent with the extension of the upstream exon using two main criteria: *i*) reads were present throughout the exon extension region in the NCLsi sample (but they can be absent from the CONTsi sample), and *ii*) normalized expression levels were consistently higher in NCLsi compared to CONTsi samples throughout the exon extension region. For further details, see Supplementary Methods.

### Gene Set Analysis

Overall, LSSs were assigned to a total of 17,975 Entrez gene IDs, with only 7,287 LSSs (0.5%) remaining unassigned. Enrichment of various biological properties among genes with activated LSSs was evaluated with the hypergeometric test against curated gene sets from the Molecular Signatures Database (Version 6.2; MSigDB) (61, 62). Additional details are available in the Supplementary Methods.

## Acknowledgments

We thank Dr. Y. Nevo for helpful discussions and suggestions, A. Petcho for excellent technical assistance, and Karoun Bagamian for editing services. This study was partially supported by grants from the Israel Science Foundation (ISF), The Israel National Center for Personalized Medicine (INCPM), and the Israel Cancer Research Fund (ICRF) to R.S., and the Helen and Milton Kimmelman Center for Biomolecular Structure and Assembly at the Weizmann Institute of Science (to J.S.). We thank The Mantoux Institute for Bioinformatics, The Crown Institute for Genomics, and The de Botton Institute for Protein Profiling of The Nancy and Stephen Grand INCPM, Weizmann Institute of Science, for mass spectrometry and analysis, and for RNA sequencing. V.G. and L.E. were supported by the Intramural Research Program of the National Human Genome Research Institute through funding granted to L.E. (1ZIAHG200323-14).

## Author Contributions

J.S. and R.S. conceived and designed the study. K.S. and A.B. performed the experiments. Y.B.C. performed the injections to *Xenopus* oocytes. R.S., V.G., and L.E. designed the RNA-Seq experiments. V.G. and L.E. designed the bioinformatic analyses. V.G. performed the bioinformatic analyses, including analysis of the RNA-Seq data. R.S., V.G., and L.E. wrote the paper. All authors discussed findings and commented on the final manuscript.

## Competing Interest Statement

The authors declare no competing interests.

## Supplementary Figure

**Figure S1.**
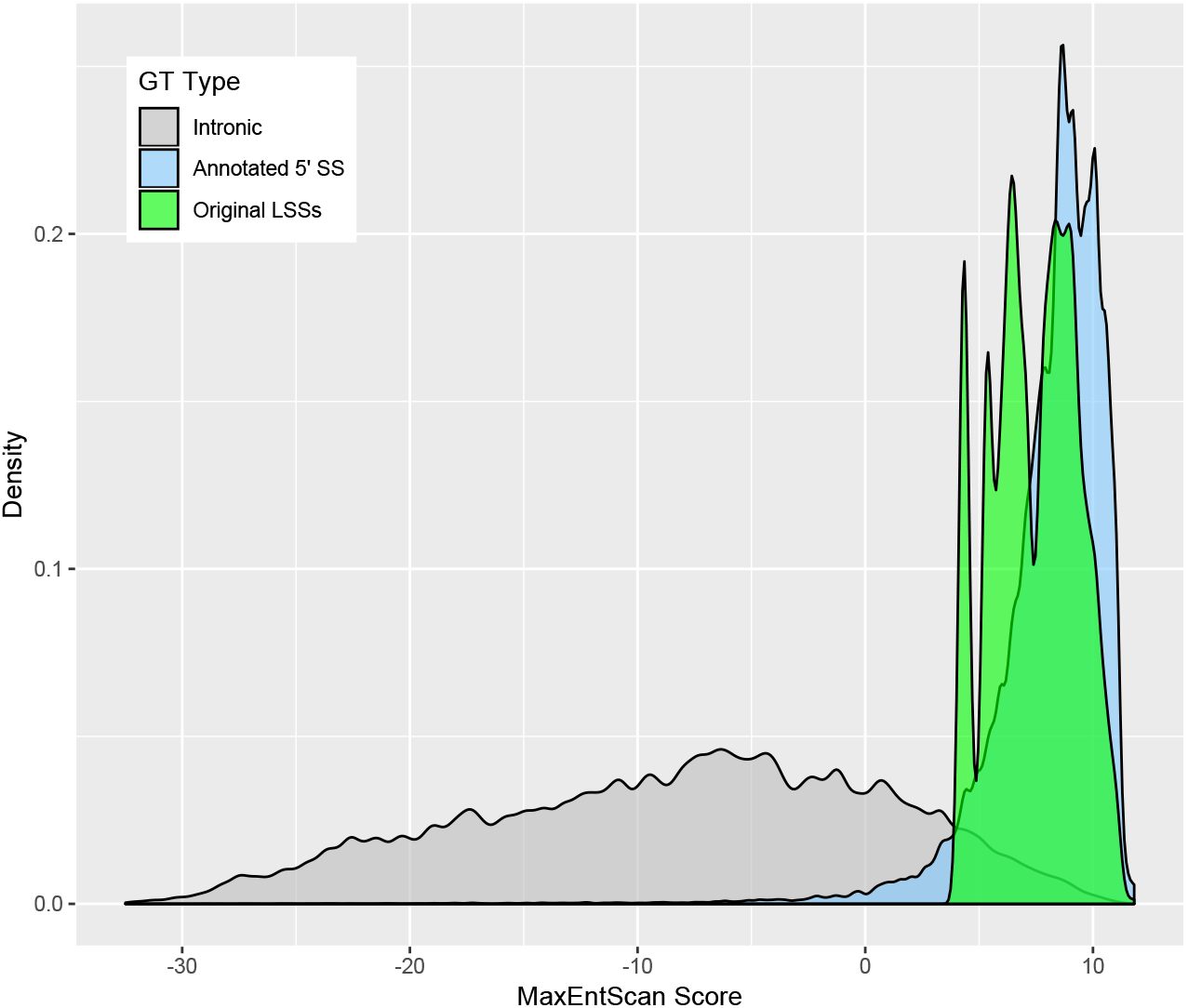
Distribution of MaxEntScan scores for GT dinucleotides in the human genome. The “Intronic” set includes 7762681 GTs located in within-CDS introns. They are limited to those GTs that would extend the upstream exon by at most 1000 nt and would allow a downstream intron of at least 20 nt. The “annotated 5’ SS” set includes 189037 GTs annotated as donor (5’) SSs in NCBI RefSeq, UCSC knownGene, or Gencode (v19) transcripts. Only within-CDS SSs were considered, and SSs with non-canonical consensus dinucleotides were excluded. The “Original LSSs” include 42305 GTs previously identified as LSSs by Nevo *et al*. (1). Their scores are based on the hg18 assembly of the human genome, whereas the other two sets are based on the data from the hg19 assembly.

## Supplementary Methods

### *In vitro* Transcription of ini-tRNA

We generated a DNA template using pTRM, a plasmid carrying the WT ini-tRNA gene (2), Q5 DNA polymerase (M0491 NEB), and specific primers in a PCR reaction according to the manufacturer’s instructions. *In vitro* transcription was carried out at 37°C for 1 hour in a reaction containing 1.5 μg DNA template, 0.5 mM NTPs, 1.5 μci a–^32^P UTP (1.8 μl BLU007H250UC, Perkin-Elmer), X1 transcription buffer, and 30 units T7 RNA polymerase (Ambion 2718). To generate biotinylated ini-tRNA, we added 0.02 mM biotin UTP to the transcription reaction. To produce cold biotinylated ini-tRNA, we added 0.06 mM cold UTP and 0.02 mM biotinylated UTP to the transcription reaction. After DNase digestion [10 U DNase (Fermentase M610A), 37°C, 20’], the RNA was purified using the miRNeasy kit (217004 Qiagen), according to the manufacturer’s instructions, and analyzed on a denaturing gel.

### Injection of ini-tRNA into the Nuclei of *Xenopus* Oocytes

We extracted *Xenopus* oocytes as previously described (3, 4). Briefly, *Xenopus* oocytes were isolated and incubated in NDE96 solution composed of ND96 (96 mM NaCl, 2 mM KCl, 1 mM CaCl_2_, 1 mM MgCl_2_, 5 mM Hepes-NaOH, pH 7.5), with the addition of 2.5 mM sodium pyruvate, 100 units/ml penicillin, and 100 μg/ml streptomycin. A day after oocyte extraction, 10 fmol (20 nl) of ^32^P ini-tRNA was injected into the oocyte’s nucleus (Picospritzer, PLI-100; Medical Systems Corp.). After 30 minutes, nuclei were extracted manually using a small needle to pierce the oocyte membrane, and the nuclei were gently squeezed out into GV buffer (5 mM HEPES, 17 mM NaCl, 83 mM KCl). Each nucleus was gently washed and transferred to a new plate (5). Radioactivity of nuclei was measured (Perkin-Elmer Tri-Carbs 2900 TR Liquid Scintillation), and only nuclei having over half of the injected ini-tRNA were further assayed. Nuclei were irradiated at 4°C with an energy of 0.8 Joule (or 0.5 Joule as indicated) [Ultra-Lum UV Cross-Linker (UVC 515)]. Nuclei were digested by RNase A [1.75X10^−3^ RNase A units (Sigma R-6513) per 50 μl sample volume] and run on 8.7% SDS PAGE (1 nucleus/well).

### RT-PCR Analysis

RT-PCR was performed on RNA extracted from the NCLsi and CONTsi treated cells as described (1), using the detailed sets of primers (see Primer List). The identity of all PCR products was confirmed by sequencing. Each experiment was repeated at least 3 times. The relative abundance was quantified in view of the intensity of GAPDH used as a control.

### Mass Spectrometry Analyses

#### Samples Preparation

We incubated 60 *Xenopus* oocyte nuclei with 120 picomol ini-tRNA (assay samples were prepared with biotinylated UTP and control samples were prepared with non-biotinylated UTP). All samples were affinity purified using 150 μl C1 streptavidin magnetic beads (as described above) and then resuspended in 100 μl 10 mM Tris buffer pH 7.5.

We used in-solution, on-bead, tryptic digestion to prepare samples. We added 8 M urea in 0.1 M Tris (pH 7.9) to the beads and incubated them (15 min at RT). Proteins were reduced by incubation with dithiothreitol (5 mM; Sigma; 60 min at RT) and alkylated with 10 mM iodoacetamide (Sigma) (30 min at RT in the dark). We diluted the urea solution to 2 M with 50 mM ammonium bicarbonate, added 250 ng trypsin (Promega; Madison, WI, USA), and incubated the reaction overnight at 37°C. We then added 100 ng trypsin for 4 hr at 37°C and stopped the digestion with 1% trifluroacetic acid (1% final concentration). Peptides were desalted using Oasis HLB μElution format (Waters, Milford, MA, USA), vacuum dried, and stored at −80°C until further analysis.

#### Liquid Chromatography

We used ULC/MS grade solvents for all chromatographic steps. Each sample was loaded using split-less nano-Ultra Performance Liquid Chromatography (10 kpsi nanoAcquity; Waters, Milford, MA, USA). The mobile phase was: A) H_2_O + 0.1% formic acid and B) acetonitrile + 0.1% formic acid. Desalting of the samples was performed online using a reversed-phase Symmetry C18 trapping column (180 μm internal diameter, 20 mm length, 5 μm particle size; Waters). The peptides were then separated using a T3 HSS nano-column (75 μm internal diameter, 250 mm length, 1.8 μm particle size; Waters) at 0.35 μL/min. Peptides were eluted from the column into the mass spectrometer using the following gradient: 4% to 30%B in 30 min, 30% to 90%B in 10 min, maintained at 90% for 5 min, and then back to initial conditions.

#### Mass Spectrometry

We coupled the nanoUPLC online through a nanoESI emitter (10 μm tip; New Objective; Woburn, MA, USA) to a quadrupole orbitrap mass spectrometer (Q Exactive Plus, Thermo Scientific) using a FlexIon nanospray apparatus (Proxeon). We acquired data in data dependent acquisition (DDA) mode, using a Top20 method. MS1 resolution was set to 70,000 (at 400m/z), mass range of 300-1650m/z, AGC of 3e6 and maximum injection time was set to 20msec. MS2 resolution was set to 17,500, quadrupole isolation 1.7m/z, AGC of 1e6, dynamic exclusion of 60sec and maximum injection time of 60msec.

#### Identification of Latent 5’SSs and Alternative Donor SSs

The list of LSSs was obtained with custom Perl software designed to scan the intronic sequences of protein-coding loci corresponding to transcripts from NCBI RefSeq (September 18, 2016), UCSC knownGene (June 30, 2013), and GENCODE v19 (December 5, 2013) annotation tracks. For this purpose we use the hg19/GRCh37 assembly of the human genome. Five criteria were used to define GTs as LSSs: *i*) they were located in introns flanked by coding exons, *ii*) they were no more than 1000 nucleotides downstream of the annotated 5’SS, and accommodating an intron of at least 20 nucleotides with the downstream 3’SS, *iii*) they scored higher than 0 on the MaxEntScan scale (6), *iv*) they were not previously annotated as the 5’SS of a separate alternative exon fully contained within the intron investigated, and *v*) the translation of the intronic region up to the LSS (i.e., the exon extension) introduced at least one premature termination codon (PTC). For LSSs contained in overlapping transcript isoforms, i.e., those with different reading frames and intron boundaries, analyses were performed separately for each isoform, and only unique LSS instances are reported for LSS counts. With the exception of the requirement for insertion of in-frame STOP codons, the same criteria were used for identifying other alternative donor splice sites (adSS_3n_, adSS_fs_). Cases of adSS_3n_ that introduce new STOP codons that span the exon junction (a total of 805 cases) were considered as adSS_fs_.

#### Analysis of RNA-Seq Data

For the genome indexing step of the STAR workflow, we included GENCODEv19 gene annotation information and information for all junctions involving LSSs and adSSs. For the alignment step, we set the “--alignSJDBoverhangMin” parameter to 2 to increase alignment sensitivity at junctions of interest, which included both previously annotated junctions and our lists of LSSs and adSSs. Overall, about 84% of reads from all four samples were uniquely mapped (**Table S3**) and used for further analyses.

#### Quantification of Increased LSS Usage

Split reads, i.e. those aligned over an intron-induced gap to exons flanking each side, can usually considered as proof of transcript undergoing the process of splicing. However, when considering cases with increased LSS usage (i.e. fold increase of at least 1.5 in the case over the control sample), we imposed additional conditions to minimize the rate of false positives due to technical artifacts. In addition to a minimum number of 4 reads required, these conditions included: *i*) the second replicate was required not to exhibit a decrease in LSS usage (i.e. increase in normalized LSS usage was require to be ≥1), *ii*) at least one split reads was required to support the LSS in the second NCLsi replicate, and *iii*) both CONTsi replicates were required to have at least 10 split reads that support the canonical junction to indicate that minimal expression can be detected for the canonical junction under normal conditions.

#### Analysis of Expression Profiles

In building the expression profiles for the exon extension regions we excluded all read pairs for which any part of either read mapped to an intronic region. In this way we minimize the impact of intronic transcriptional noise on evaluating the expression profiles, which is variable across the human genome. Additionally, in evaluating the expression profile, we allowed small deviations from the condition of consistent higher expression level in the NCLsi sample in order to accommodate situation in which reads not originating from the transcripts of interest (e.g. reads generated by intronic transcriptional noise that fully map to exonic regions, reads that originate from alternative isoforms) cannot be readily eliminated. Our pipeline allowed us to eliminate many cases inconsistent with activation of latent splicing at LSS but which showed an increase in LSS usage. Such cases included, but are not limited to, unannotated exons, intronic alternative transcription start sites, or specific amplification of split reads that can happen during construction of sequencing library.

#### Gene Sets Analysis

Assignment of LSS-containing transcripts to Entrez gene IDs was performed using six cross-reference datasets: “gene2refseq” and “gene2ensembl” from NCBI (ftp://ftp.ncbi.nlm.nih.gov/gene/DATA/, accessed on October 30, 2018), “knownToRefSeq and “knownToEnsembl” from UCSC (http://hgdownload.soe.ucsc.edu/goldenPath/hg19/database/, accessed on October 30, 2018), the GENCODE v19 annotation set, as well as a custom reference dataset downloaded through the BioMart interface (https://www.ensembl.org/biomart, accessed on October 30, 2018) containing: Ensembl gene ID, Ensembl transcript ID, NCBI RefSeq and UCSC transcript accession numbers, gene symbol, and NCBI Entrez gene ID. The sets of curate genes were obtained from version 6.2 of MSigDB, which contains 8 gene set collections. Each collection contains between 50 and 5917 individual gene sets (17,810 gene sets in total). Each gene set consists of between 5 and 2940 unique gene identifiers, provided as gene symbols and Entrez gene IDs. To correctly evaluate enrichment in biological properties for the genes with activated LSSs, all genes included in the analysis (i.e. genes from the MSigDB gene sets, as well as genes in the background set) were required to have evidence of *i*) at least one LSS with a MaxEntScan score above 0.05 and *ii*) support of at least 10 split reads in both CONTsi samples for the canonical junction encompassing the suitable LSSs in order to match the genes that could be detected as having activated LSSs.

### Primer List

Specific primers for **ini-tRNA** generation were designed:
Sense primer: 5’-TAATACGACTCACTATAAGCAG - 3’,
and antisense primer: 5’ - TGGTAGCAGAGGATGGTTTC-3’

**elongator ini-tRNA** template primer:
5’TGGTGCCCCGTGTGAGGATCGAACTCACGACCTTCAGATTATGAGACTGACGCGCTACCTA
CTGCGCTAACGAGGCTATAGTGAGTCGTATTA-3
T7 primer: 5’-TAATACGACTCACTATA-3’

Primer sets for the RT-PCR validation experiments:

**RAF1**
Sense primer 5’-TCTGGCATGTTGAGGGCTTTG - 3’
Antisense primer 5’-ACTTTGGTGCTACAGTGCTCA - 3’
Annealing temp 57° C

**ANK2**
Sense primer 5’-GAGCTTGTGTCCTCTACATG - 3’
Antisense primer 5’-TCCAGCTGTCCCCATTCTCA - 3’
Annealing temp 56° C

**VRK2**
Sense primer 5’-GCAAAGCAAGCTACTTCCAT - 3’
Antisense primer 5’-AATTCAGTCAGACCAGATCC - 3’
Annealing temp 51.5°C

**KIAA0169**
Sense primer 5’-TGTGGCACAGCCCCTATTCCA - 3’
Antisense primer 5’-AGACCAAAGGTTCCCAAGGTG - 3’
Annealing temp 58° C

**COPA**
Sense primer 5’-CAGAAGCGGTAACATATCAGG - 3’
Antisense primer 5’-CGTGTTTGGCTAGTAGTGCT - 3’
Annealing temp 51°C

**LARS**
Sense primer 5’-TTGCGGCAGAAAATAGACCTA - 3’
Antisense primer 5’-GTCACACAGAGCCACAACAC - 3
Annealing temp 55° C

**HTT**
Sense primer 5’-CAGGTCGTCCTTGTGTTAGGA - 3’
Antisense primer 5 - CTTGCATGGTGGAGAGACGA - 3’
Annealing temp 57° C

**TCERG1**
Sense primer 5’-GTCAGAAAACAATCTTTGGGGG - 3’
Antisense primer 5’-TGTGCAACTCCTTCTCCCAC - 3’
Annealing temp 55° C

**HEATR1**
Sense primer 5’-CAATTGAGAAATGGTGCTAGTC - 3’
Antisense primer 5’-TGATGAATGATGGAGACGACCA - 3’
Annealing temp 5l.5°C

**GAPDH**
Sense primer 5’-CCAGCCGAGCCACATCGCTC - 3’
Antisense primer 5’ - TGAGCCCCAGCCTTCTCCAT - 3’
Annealing temp 60.0°C

## Supplementary Tables

**Table S1.** List of peptides identified by mass spectrometry.

**Table S2.** List of latent splice sites (LSSs) identified in the human genome.

**Table S3.** Summary of reads obtained through the RNA-Seq experiment and mapped to each of the four samples.

**Table S4.** List of genes with differential expression in NCLsi compared to CONTsi samples.

**Table S5.** List of LSSs activated in NCLsi samples upon NCL knockdown.

**Table S6.** Gene sets over-represented among the genes with activated LSSs.

**Table S7.** List of LSSs activated in CONTsi samples.

**Table S8.** List of adSS_3n_ splice sites identified in the human genome.

**Table S9.** List of adSS_3n_ activated in NCLsi samples upon NCL knockdown.

**Table S10.** List of adSS_3n_ activated in CONTsi samples.

**Table S11.** List of adSS_fs_ splice sites identified in the human genome.

**Table S12.** List of adSS_fs_ activated in NCLsi samples upon NCL knockdown.

**Table S13.** List of adSS_fs_ activated in CONTsi samples.

## Notes

### Competing Interest Statement

The authors have declared no competing interest.

